# Making a pathogen? Evaluating the impact of protist predation on the evolution of virulence in *Serratia marcescens*

**DOI:** 10.1101/2022.06.17.496625

**Authors:** Heather A. Hopkins, Christian Lopezguerra, Meng-Jia Lau, Kasie Raymann

## Abstract

Opportunistic pathogens are environmental microbes that are generally harmless and only occasionally cause disease. Unlike obligate pathogens, the growth and survival of opportunistic pathogens does not rely on host infection or transmission. Their versatile lifestyles make it challenging to decipher how and why virulence has evolved in opportunistic pathogens. The Coincidental Evolution Hypothesis (CEH) postulates that virulence results from exaptation or pleiotropy, i.e., traits evolved for adaptation to living in one environment that have a different function in another. In particular, adaptation to avoid or survive protist predation has been suggested to contribute to the evolution of bacterial virulence (the training grounds hypothesis). Here we used experimental evolution to determine how the selective pressure imposed by a protist predator impacts the virulence and fitness of a ubiquitous environmental opportunistic bacterial pathogen that has acquired multi-drug resistance: *Serratia marcescens*. To this aim, we evolved *S. marcescens* in the presence or absence of generalist protist predator, *Tetrahymena thermophila*. After 60 days of evolution, we evaluated genotypic and phenotypic changes by comparing evolved *S. marcescens* to the ancestral strain. Whole genome shotgun (WGS) sequencing of the entire evolved populations and individual isolates revealed numerous cases of parallel evolution, many more than statistically expected by chance, in genes associated with virulence. Our phenotypic assays suggested that evolution in the presence of a predator maintained virulence, whereas evolution in the absence of a predator resulted in attenuated virulence. We also found a significant correlation between virulence, biofilm formation, and grazing resistance. Overall, our results provide evidence that bacterial virulence and virulence related traits are maintained by selective pressures imposed by protist predation.

## Introduction

Opportunistic pathogens are environmental bacteria that typically coexist peacefully alongside hosts and only sometimes cause disease, e.g., in hosts with immunodeficiency or microbiome imbalance (1). Opportunistic pathogens are intriguing because they have diverse lifestyles and experience strong tradeoffs between adaptation to life in the environment versus life inside of a host (1–3). Opportunistic pathogens are also conceptually challenging because, unlike obligate pathogens, their survival does not depend on their ability to infect a host: they are fully capable of reproducing and living in the environment (1, 4). Thus, the factors driving virulence in opportunistic pathogens remain largely enigmatic from an evolutionary standpoint. Several hypotheses have been put forth, proposing that virulence is selected for, or maintained by, interactions that occur outside a host. The Coincidental Evolution Hypothesis (CEH) postulates that virulence evolves indirectly due to selection that occurs in non-host environments (5, 6), and protists have been proposed to act as “Trojan Horses” (7) or training grounds (8) for the evolution of bacterial pathogens. These hypotheses stem from the fact that bacterial defense mechanisms against environmental predators (e.g., protists) such as adherence, biofilm formation, grazing resistance, intracellular toxin production, and outer-membrane structures also play a role in host infection (1, 9, 10). Additionally, several studies have shown that pathogens challenged with microbial predators exhibited increased host virulence (11–16).

Experimental evolution can help reveal how different environmental pressures impact the evolution of bacteria and which genes are involved in specific phenotypic traits (17–19). In recent years, experimental evolution has been used to investigate if eukaryotic predation increases the virulence of opportunistic bacterial pathogens, but results have been conflicting (20–26). One major limitation of many previous studies has been the lack of thorough phenotypic and genotypic characterization. Not all populations will follow the same evolutionary trajectory (27–31). Therefore, evaluating and comparing genotype and multiple phenotypes in experimental evolution studies is critical. For example, even if predation generally selects for the maintenance or enhancement of pathogen virulence, it is possible that some traits that are adaptive for defense against predators have no impact on virulence or even result in attenuation of pathogenicity within hosts. Nevertheless, experimental evolution can be used to answer many questions about fundamental evolutionary processes (17–19) and help reveal genes and mechanisms associated with virulence and other phenotypes. Thus, the goal of this study was to experimentally evolve the opportunistic bacterial pathogen *Serratia marcescens* in the presence or absence of an environmental protist predator and characterize both the genotypes and phenotypes of the evolved bacteria to elucidate how predation, or the lack thereof, impacts opportunistic pathogen evolution. We hypothesized that virulence would be increased or at least maintained in the presence of a predator.

*S. marcescens* is an opportunistic pathogen that ubiquitously occurs in soil and water (32) and has been used for several bacteria-predator experimental evolution studies (20, 21, 23). It is a host generalist that can colonize many different organisms, where it can be pathogenic and/or commensal (32). In humans, *S. marcescens* is a common enteric bacterium that is generally harmless in the gastrointestinal tract (33). However, it was recently found to be capable of damaging gut epithelial cells *in vitro* (34) and is responsible for causing dangerous nosocomial infections worldwide, notably in the respiratory and urinary tracts (35, 36). In the last few decades, the frequency of nosocomial infections caused by *S. marcescens* has increased, several hospital-wide outbreaks have been reported, and most strains have acquired multidrug resistance (35–37). Additionally, *S. marcescens* has been recognized as an opportunistic pathogen of many plants and other animals (32). Thus, understanding the evolution of virulence in *S. marcescens* is important for the health of a wide range of organisms. In this study, we used *S. marcescens* KZ19 (38), which was isolated from the gut of a honey bee (*Apis mellifera*) and is pathogenic to honey bees following microbiome disruption (39–42) or if administered orally to bees at high doses (38). This strain of *S. marcescens* resides in a monophyletic clade with human and plant isolates, with the closest sequenced relative being a nosocomial strain BIDMC 50 (38).

Here, we serially passaged *S. marcescens* KZ19 every day for 60 days in the presence or absence of a generalist protist predator, *Tetrahymena thermophila* (Figure 1). Following 60 days of evolution, we performed whole genome shotgun (WGS) metagenomic sequencing on the entire *S. marcescens* evolved populations, sequenced individual evolved isolates, and characterized the mutations present in both evolved isolates and populations. We observed multiple cases of parallel evolution which, when combined with phenotypic assays of isolates, enabled us to predict genes and pathways involved in virulence-related phenotypes. Although all populations did not follow the same evolutionary trajectories, in general, we found that virulence and grazing resistance was attenuated in the absence of a predator and predation resulted in increased biofilm production and slightly elevated virulence. Additionally, we found strong correlations between predation (grazing) resistance, biofilm production, and pathogenicity (in honey bees), lending support to the hypothesis that predation resistance and virulence are often governed by the same mechanisms. In conclusion, our study provides evidence that experimental evolution can be used to help elucidate how exposure to non-host environments impacts opportunistic pathogen evolution.

**Figure 1.**
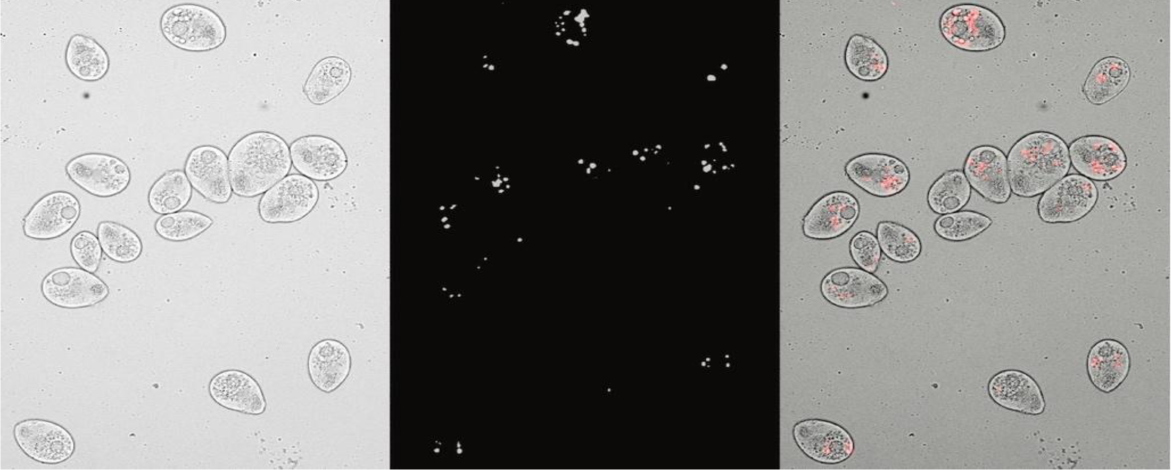
Confirmation of grazing activity. A) Phase contrast image of *T. thermophila* after 24h co-culture with fluorescently labeled *S. marcescens* KZ19 in Neff media with 180 ug/ml spectinomycin, B) Cy5 filter image showing the E2 crimson labeled *S. marcescens* KZ19 cells inside *T. thermophila*, and C) composite overlay of the phase contrast and fluorescent images. Images were taken at 20X on a Keyence BZ-X700 Series All-In-One Fluorescence Microscope and overlayed in ImageJ.

## Results

We confirmed that *T. thermophila* grazes on the ancestral bacterial strain by visualizing a 24h co-culture of *T. thermophila* and a transconjugant *S. marcescens* KZ19 containing an E2 crimson fluorescent protein (38, 43) via fluorescent microscopy (Figure 1). We then created six media-evolved (ME) and six predator-evolved (PE) replicate Lines (populations). We minimized predator/prey co-evolution by passaging evolved *S. marcescens* to new axenic *T. thermophila* cultures each day. The same concentration of *S. marcescens* was transferred for each passage, and we confirmed successful passaging via plating on LB agar every day. Viability and population size of *T. thermophila* was evaluated before every passage via light microscopy. Following 60 days of evolution the entire PE (*n*=6) and ME populations (*n*=6) were sequenced using WGS metagenomic sequencing. The ancestor (*n*=1) and individual isolates from the three ME (Lines 1-3) and six PE (Lines 1-6) populations were sequenced using WGS sequencing (Table S1). We then performed phenotypic assays on the evolved isolates (ME=3 and PE=6) in order to associate mutations with phenotypes.

### Mutations in the predator- and media-evolved populations

All evolved populations were compared to the ancestral genome to identify mutations. First, the ancestral genome was re-sequenced using both long (Nanopore) and short (Illumina) read sequencing technologies and assembled using SPades v3.15.3 (44) to generate a consensus ancestral genome (Table S1). We then corrected for false positives (i.e., sequencing errors) by mapping all the short reads (244X coverage) of the ancestral strain back to the new consensus ancestral genome using BreSeq v0.38.2 (45). All mutations predicted via BreSeq (45) when the ancestral reads were mapped to the ancestral consensus genome were considered false positives if detected in the evolved populations (i.e., they most likely correspond to sequencing or assembly issues in the ancestral genome).

We first analyzed the mutation patterns across each population. The total number of mutations identified in each evolved population (ME_p_ *n*=6; PE_p_ *n*=6) ranged from 6 to 106 (minimum frequency cutoff of 0.05) with an average of 44 mutations per population (Supplemental Dataset 1). The population with the lowest mutations identified was ME_p_3 and the two with the largest number of mutations were PE_p_3 and ME_p_2. However, the number of mutations present in a population was negatively correlated to the average genome coverage (*P*= 0.0001, *R^2^=*0.7805; Figure S1). A negative correlation between the number of mutations and genome coverage indicates that our sequencing depth was sufficient to characterize the mutations present in the populations.

In all evolved populations we identified nonsynonymous, synonymous, and intergenic mutations (Figure 2A, Supplemental Dataset 1). To determine the percentage of nonsynonymous, synonymous, intergenic, non-sense, and RNA mutations expected at random, we simulated independent mutations in the ancestral *S. marcescens* KZ19 genome by performing 10,000 independent simulations using a Jukes & Cantor model (47). We found that the percentage of nonsynonymous mutations detected in our populations (49-68%) was between 4-23% less than the random expectation of 71.5% (Figure 2A), indicating that overall, both environments imposed negative (purifying) selective pressure on *S. marcescens* (i.e., deleterious mutations are being purged).

**Figure 2.**
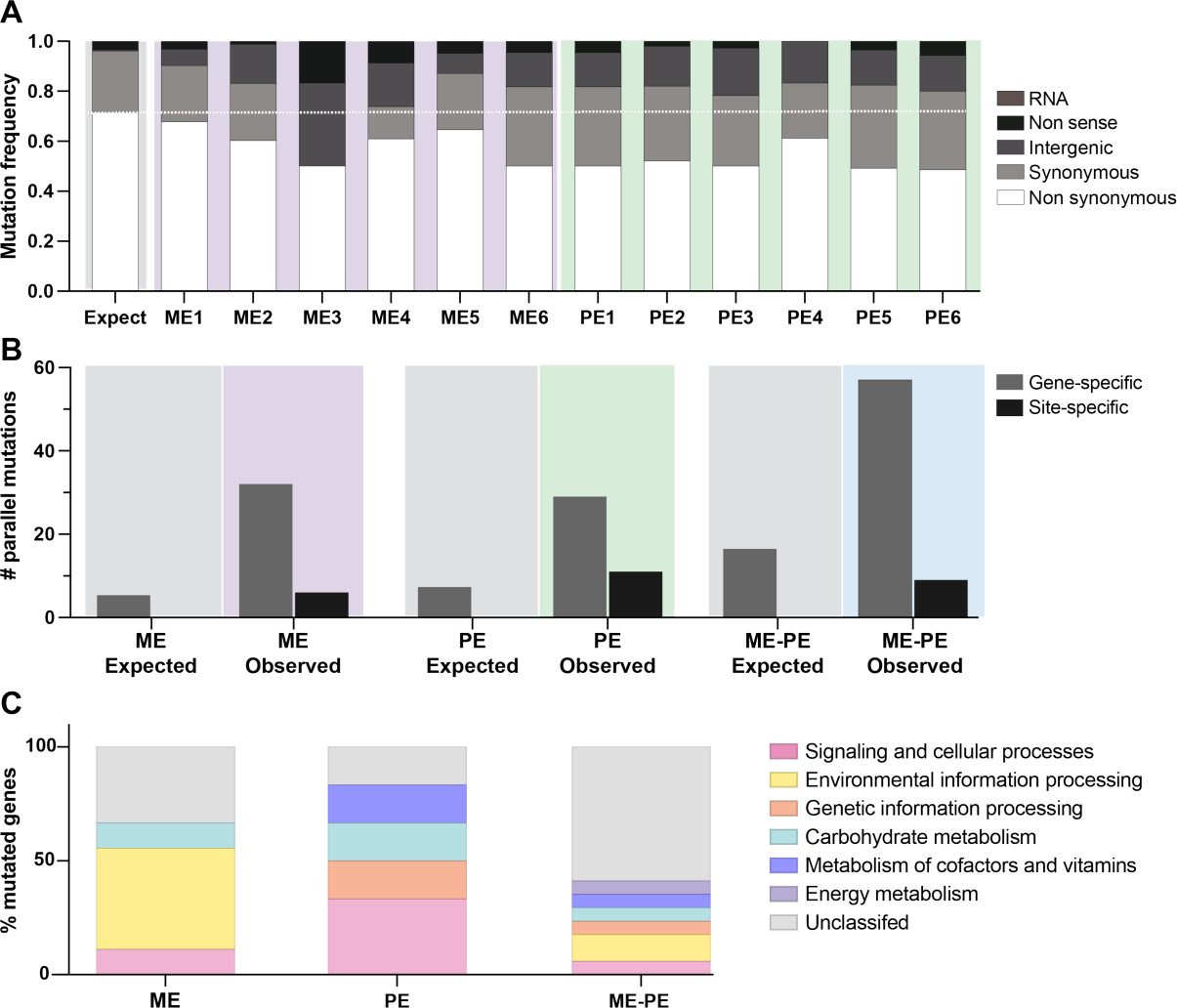
Characterization of the mutations present in the evolved populations. **A)** Percentage of nonsynonymous, synonymous, intergenic, non-sense, and RNA mutations detected in each evolved population (ME1-6 and PE1-6) and compared to the percentage of each mutation type expected at random (Expected). **B)** Number of observed parallel (*gene-* and *site-specific*) mutation events observed in ME, PE, and in both ME and PE (ME-PE) populations compared to random expectation (expected vs. observed). Significantly more *gene-specific* and *site-specific* parallel mutations were observed than would be expected to occur by chance (Chi-squared test, *P*<0.0001). **C)** the KEGG (46) functional annotations of the genes in which parallel events occurred.

In addition to purifying selection, we also found evidence that our populations were undergoing positive selection based on the identification of numerous mutations either in the exact same position (*site-specific* parallel evolution) or in the same gene (*gene-specific* parallel evolution) across populations (Figure 2B, Supplemental Dataset 1). Parallel evolution events were considered if a mutation was found in the same position or gene in independently evolved populations (ME_p_, PE_p_, or both). None of the parallel evolution events occurred across all populations, confirming that they were not present in the ancestral population. We observed a total of 118 parallel mutations; 57 (nine *same-site*) were shared between the ME_p_ and PE_p_ populations, 29 (11 *same-site*) were unique to PE_p_ populations, and 32 (six *same-site*) were unique to ME_p_ populations (Figure 2B, Supplemental Dataset 1).

To determine the number of parallel mutations expected to occur by chance we simulated independent mutations in the ancestral *S. marcescens* KZ19 genome by performing 10,000 independent simulations using a Jukes & Cantor model (47). The number of *site-specific* parallel mutations expected to occur by chance within or across all our evolved populations was less than one, indicating that we found overwhelmingly more site-specific parallel mutations than random expectation (Figure 2B, ME_p_ *P*=2.2e-16, PE_p_ *P*= 2.2e-16, ME_p_-PE_p_ P= 2.2e-16, Chi-squared test). The number of *gene-specific* parallel mutations observed across the ME_p_ and PE_p_ populations was also four times more frequent than expected at random (Figure 2B, ME_p_ *P*= 8.7e-25, PE_p_ *P*= 6.2e-05, ME_p_-PE_p_ P= 6.4e-10, Chi-squared test). The presence of excessive convergent parallel mutations provides strong evidence that positive selection occurred in all populations to adapt to their respective environments (the media and the presence of a predator).

Parallel evolution events occurred in 32 different genes and eight different promoter regions. Of the 32 genes in which we identified parallel evolution events, 70% could be functionally classified (Figure 2C). Although parallel evolution events arose across and within predator (PE_p_) and media (ME_p_) evolved populations, both Lines were impacted in different types of genes. A large portion of the parallel mutations that occurred across only ME_p_ populations were in genes involved in environmental information processing, while the parallel evolution events unique to PE populations were concentrated in genes involved in signaling and cellular processes. Overall, parallel mutations were identified in genes involved in genetic information and processing (PE_p_=17%, ME_p_=0%, PE_p_+ME_p_=6%), signaling and cellular processes (PE_p_=33%, ME_p_=11%, PE_p_+ME_p_=6%), environmental information processing (PE_p_=0%, ME_p_=44%, PE_p_+ME_p_=12%), and metabolism (PE_p_=33%, ME_p_=11%, and PE_p_+ME_p_=18%), and unclassified genes (PE_p_=17%, ME_p_=33%, PE_p_+ME_p_=59%). Parallel mutations present in only ME or shared between PE and ME populations are likely conferring adaptation to the Neff media, whereas parallel mutations only present in the PE populations suggest that they are important for fitness in the presence of a predator. Since most of the genes in which we identified mutations have not been functionally confirmed in *S. marcescens*, all gene names and predicted functions discussed hereafter must be considered candidate genes/functions.

We plotted all the mutations identified in each ME_p_ and PE_p_ population and their observed frequencies and found that several parallel mutations were fixed within the populations (Figure 3A). Interestingly, most of the fixed parallel mutations occurred in genes within the same protein-protein interaction network predicted via STRING v12 (48) in close relatives of *S. marcescens*, such as *S. rubidaea* and *Yersina enterocolitica* (Figure 3B). This protein network contains the BarA-UvrY two component regulatory system and the three core proteins of the Rcs phosphorelay system, (*rcsC*, *rcsD*, and *rcsB*). Both BarA-UvrY and the Rcs regulatory systems have been shown to play a major role in environmental adaptation and the regulation of virulence in *S. marcescens* (49–51) or related enterobacteria (52–56). Mutations in the histidine kinase *barA* were found in multiple PE_p_ (3/6) and ME_p_ populations (5/6), whereas mutations in the response regulator *uvrY* were mostly restricted to ME_p_ populations (Figure 3C). Mutations in the response regulator of the Rcs system, *rcsB*, were identified at high frequency in virtually all ME_p_ populations (5/6) but never in the PE_p_ populations (Figure 3C). One ME_p_ population (ME_p_6) contained a fixed mutation in the transmembrane hybrid kinase *rcsC* and one PE_p_ population (PE_p_3) also had a mutation in *rcsC* but at very low (<10%) frequency (Figure 3C). Within this network, fixed (100 percent frequency) *site-specific* parallel mutations unique to ME_p_ populations (ME_p_2-6) were identified in a promoter region upstream of a gene annotated as the biofilm dispersion protein *bdlA* (Figure 3C). Four of the six PE_p_ populations (PE_p_2, PE_p_4, PE_p_5, and PE_p_6) contained mutations in a gene annotated as the diguanylate cyclase *yfiN*, and of these four populations, three (PE_p_4-6) also contained mutations in a promoter upstream of a gene annotated as the outer membrane protein *ompC* (Figure 3C). The genes *bdlA*, *yfiN*, and *ompC* have all been implicated in biofilm formation in other *Enterobacteriaceae* species (57–62). It is important to note that the six PE_p_ (Lines 1-6) and three ME_p_ (Lines 1-3) populations were evolved at the same time, but three of the ME_p_ (Lines 4-6) populations were started over one year later, demonstrating the reproducibility of our results.

**Figure 3.**
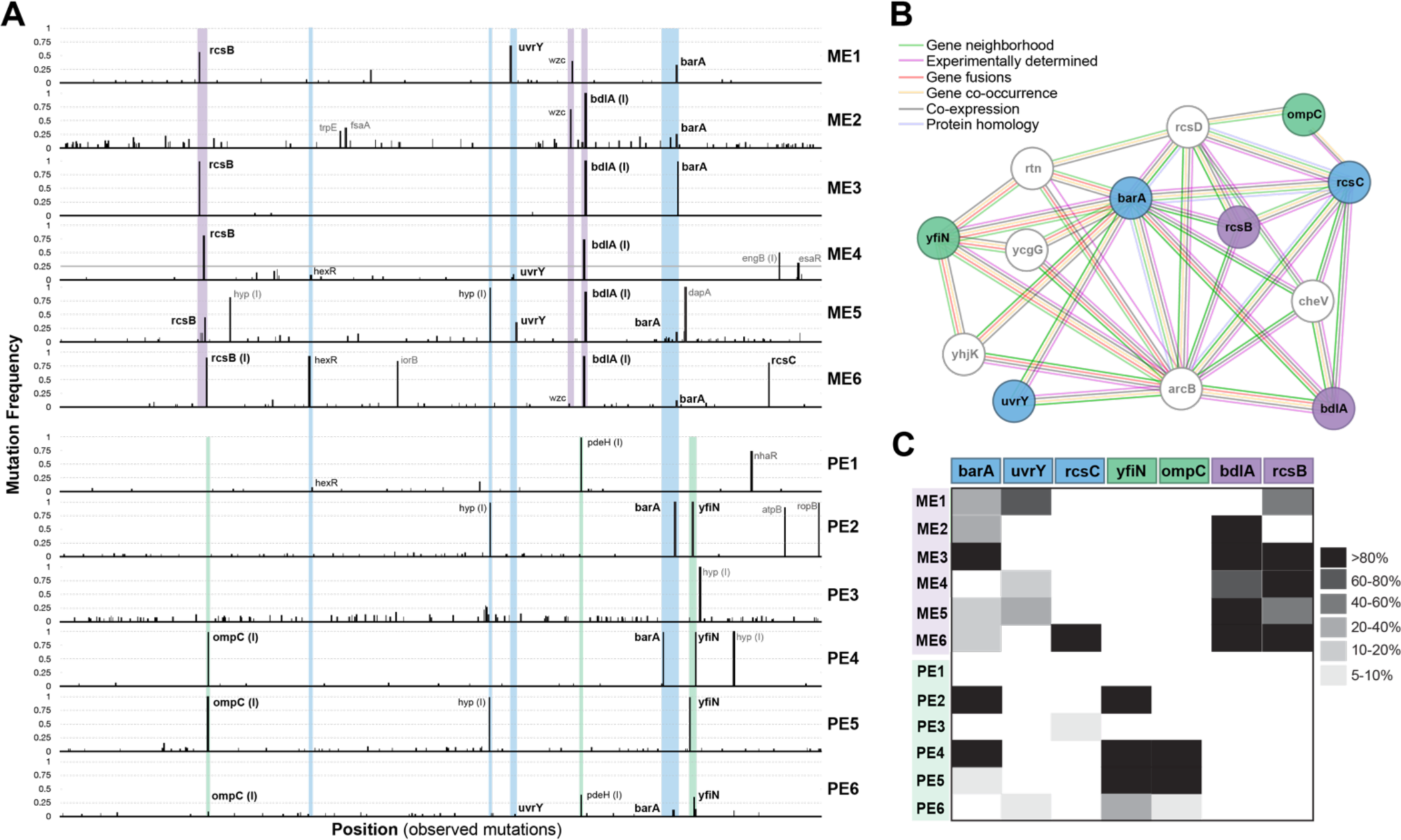
Frequency of mutations present in the evolved populations. **A)** Plot of all the detected mutations and their frequencies in each population highlighting the parallel and/or “fixed” mutations. Cases of parallel mutation events are highlighted in green (PE only mutations), purple (ME only mutations), and blue (ME-PE). **B)** STRING v12 (48) predicted protein-protein interaction network containing seven genes in which we identified multiple parallel mutations. **C)** Heatmap showing the frequency of mutations in or adjacent (indicated by *) to the seven genes from the same protein-interaction network within each evolved population.

### Genotype and phenotype of predator- and media-evolved isolates

To attempt to link specific mutations with phenotypes, a total of three ME_i_ and six PE_i_ isolates were sequenced from the evolved populations and compared to the ancestral genome to identify mutations (Figure 4A, Table S2). At least one mutation was detected in each isolate, with a maximum of four and an average of two mutations per isolate (Figure 4A). We identified a total of 24 mutations within or in upstream promoters of 14 genes and 70% of these mutations (17/24) were present at a frequency of >90% in their respective population (Table S2). Additionally, the isolates possessed all the fixed (100% frequency) mutations present within the population from which they were isolated (Figure 3A, Supplemental Dataset 1).

**Figure 4.**
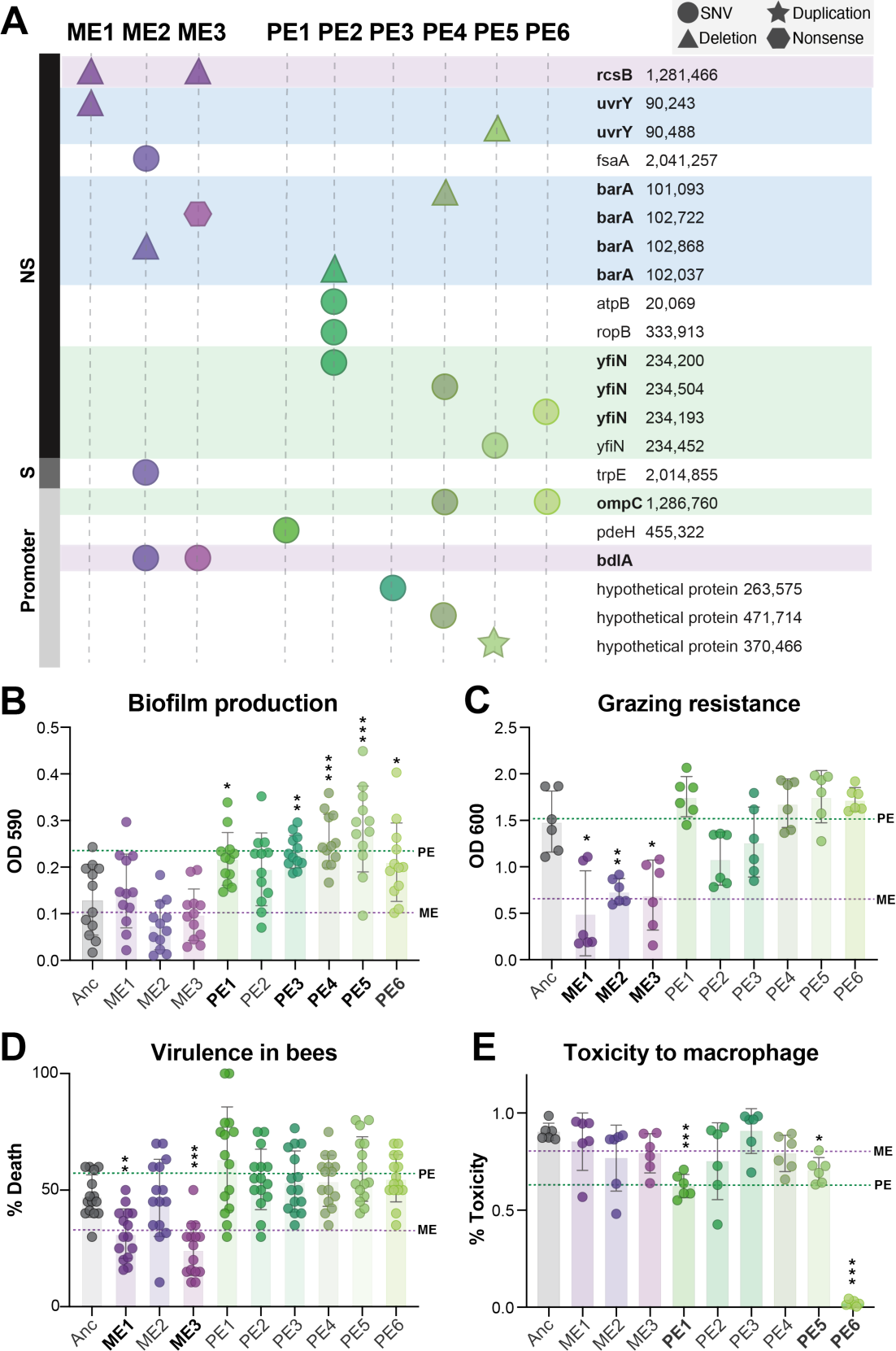
Genotypes and phenotypes of evolved isolates. **A)** Mutations present in each isolate grouped by nonsynonymous (NS), synonymous (S), promoter (intergenic mutation in a promotor region) and indicating the mutation type (single nucleotide variant (SNV), nonsense mutation, duplication, or deletion) and the gene (or gene upstream) affected. **B)** Mean biofilm formation of MEi and PEi isolates and the ancestral strain based on 590nm OD readings after 24h of growth in Neff media. Four assays were performed in triplicate and each point represents a biological replicate. **C)** Mean grazing (predation) resistance of PEi and MEi isolates and the ancestral strain based on 600nm OD readings of bacterial filtrate after 48h in co-culture with *T. thermophila*. Two assays were performed in triplicate and each point represents a biological replicate. **D)** Percent death of honey bees five days after exposure to the ancestral, MEi, and PEi isolates (control group average percent death = 2.3%; see Figure S4). Three replicate assays were performed with five replicates of 20 bees each per isolate per assay (15 replicates total per isolate). Each data point represents a biological replicate of 20 bees. **E)** Percent cytotoxicity of the ancestor, MEi, and PEi isolates to murine macrophages (RAW264.7) based on LDH release. **B-E)** Each data point represents a biological replicate (n=6). All box plots show the mean standard deviation. Significance was tested by comparing each evolved isolate to the others and the ancestor using one-way ANOVA with Dunnett’s multiple comparisons test. * = P<0.01, **= P<0.001, ***= P<0.0001. Dashed lines represent the overall mean when combining all MEi (purple line) or PEi (green line) isolates, respectively.

We identified three instances of *site-specific* parallel evolution and three cases of *gene-specific* parallel evolution across the isolates (Figure 4A), all of which occurred within coding regions or promoters upstream of genes within the same BarA-UvrY Rcs protein-protein interaction network (Figure 3B). Two of the three *site-specific* parallel mutations were unique to ME_i_ isolates: a 66bp deletion in rcsB (ME_i_1 and ME_i_3) and an intergenic SNV in a promoter upstream of a gene annotated as bdlA (ME_i_2 and ME_i_3; Figure 4A). The third *site-specific* mutation was unique to PE_i_ isolates: an intergenic SNV in a promoter upstream of *ompC* (PE_i_4 and PE_i_6; Figure 4A). Two of the *gene-specific* parallel evolution events occurred in both ME and PE isolates: *barA* (ME_i_2, MEv3, PEv2, and PEv4) and *uvrY* (ME_i_1 and PE_i_5). The third case of *gene-specific* parallel evolution arose in a gene annotated as the inner membrane protein *yfiN* in all PE_i_ isolates except PE_i_1 and PE_i_3 (Figure 4A). Overall, most of the mutations present in the isolates likely represent adaptive mutations as they were fixed in their respective populations.

To evaluate the phenotypes of the PE_i_ and ME_i_ isolates, we performed growth, biofilm production, predation resistance (population size after growth in the presence of *T. thermophila*), and virulence (mortality of honey bees and death of murine macrophage) assays. We chose to specifically analyze isolates, rather than populations, in order to correlate fixed mutations with phenotypic characteristics. To compare growth between the ancestral and evolved strains, we monitored optical density (OD) in spent Neff media (filtered media that had supported *T. thermophila* growth for 48h), fresh Neff media, and LB media every hour for 24 hours. None of the ME_i_ and PE_i_ isolates significantly differed in growth in either media type when compared to the ancestor (Figure S2). Because biofilm formation is a strategy for resistance to predation and is also associated with virulence, we measured biofilm production in our ME_i_ and PE_i_ isolates compared to the ancestor (Figure 4B). None of the ME_i_ isolates significantly differed in biofilm production when compared to the ancestor, whereas five of the six PE isolates produced significantly more biofilms than the ancestor (Figure 4B; PE_i_1 *P*=0.017, PE_i_3 *P*=0.003, PE_i_4 *P*=0.0001, PE_i_5 *P*<0.0001, PE_i_6 *P*=0.035, one-way ANOVA with Dunnett’s multiple comparison test). Overall, when considered together, mean biofilm production of the PE_i_ isolates was significantly higher than the mean of the ME_i_ isolates and the ancestor (Figure S3A, P<0.01, with one-way ANOVA with Dunnett’s multiple comparison test).

In order to determine if exposure to a predator resulted in increased predation resistance, we evaluated grazing resistance (i.e., ability to proliferate in co-culture with a predator) of the ME_i_ and PE_i_ isolates. Each isolate was grown with *T. thermophila* for 48h and then the population density of the bacteria was measured. All ME_i_ isolates were found to be significantly more susceptible to predation than the ancestor (Figure 4C; *P<* 0.0001, one-way ANOVA with Dunnett’s multiple comparison test). In contrast, none of the PE_i_ isolates significantly differed in resistance to predation relative to the ancestor. However, the mean grazing resistance of all PEi isolates combined was significantly higher than the mean of the ME_i_ isolates (Figure S3B, P<0.01, with one-way ANOVA with Dunnett’s multiple comparison test).

We further determined how evolution under predation pressure impacts the virulence of *S. marcescens* KZ19 by performing survival assays in a natural host, the honey bee. Honey bees were orally exposed to each individual isolate, the ancestral strain, or sucrose solution only (controls) and monitored every day for five days (Figure S3). We evaluated the percentage of death, five days post-exposure and found that two of the three ME isolates (ME_i_1 and ME_i_3) displayed significantly attenuated virulence when compared to the ancestral strain (Figure 4D; ME1 *P*=0.001, ME3 *P*<0.0001, one-way ANOVA with Dunnett’s multiple comparison test). Conversely, none of the PE_i_ isolates significantly differed from the ancestor in terms of virulence (Figure 4D). Based on probability of survival using a Kaplan-Meier method, we obtained the same results, with only ME_i_1 and ME_i_3 displaying significantly decreased virulence when compared to the ancestor (Figure S4, P<0.05, Mantel-Cox Log-rank test with Bonferroni correction). However, when comparing the mean percent death five days post-exposure by combining all ME_i_ and PE_i_ isolates, the ME_i_ isolates displayed significant attenuation of virulence when compared to the ancestor and PE_i_ isolates, and the PE_i_ isolates were significantly more virulent than the ancestor and the ME_i_ isolates (Figure 3SC, P<0.01, with one-way ANOVA with Dunnett’s multiple comparison test).

Several clinical *S. marcescens* isolates have been shown to be cytotoxic to macrophages (63–65). Although *S. marcescens* KZ19 was isolated from the gut of a honey bee, we found that it is capable of killing RAW 264.7 murine macrophages (Figure 4E). Thus, we evaluated the percent of murine macrophage killing by the ancestor and compared it to the ME_i_ and PE_i_ isolates based on the release of lactate dehydrogenase (LDH) after two hours of co-culture (MOI 10:1). None of the ME_i_ isolates differed from the ancestor in terms of cytotoxicity to macrophages. Conversely, three of the PE_i_ isolates displayed significantly less cytotoxicity to macrophage when compared to the ancestral strain (Figure 4E; PE_i_1 *P*= 0.0009, PE_i_5 *P*=0.03, PE_i_6 *P*<0.0001, one-way ANOVA with Dunnett’s multiple comparison test). When all ME_i_ and PE_i_ isolates were combined, no difference was observed between the ME_i_ isolates and the ancestor or the PE_i_ and ME_i_ isolates, but the PE_i_ isolates were significantly less cytotoxic to macrophage cells than the ancestor (Figure 3SD). However, macrophage killing appears to be time and density dependent. In a separate timeseries experiment with a different starting ratio (MOI 1:10), we found that all evolved isolates (ME_i_ and PE_i_) displayed overall less cytotoxicity to macrophage than the ancestor at hours 2-5 post exposure, but after six hours they converged in their level of cytotoxicity with all macrophages being killed (lysed) by all isolates and the ancestral strain (Figure S4). Therefore, macrophage cytotoxicity may not be an informative metric for gauging virulence in *S. marcescens* KZ19.

Finally, we tested whether there was a direct link between pathogenicity in honey bees, cytotoxicity to murine macrophage, biofilm production, and grazing resistance (Figure 5). Despite our relatively small sample size, we found a significant positive correlation between biofilm production and grazing resistance (Figure 5A; *r^2^*=0.56, *P*=0.01, Pearson correlation) as well as between virulence and biofilm production (Figure 5B; *r^2^*=0.5, *P*=0.02, Pearson correlation) and virulence and grazing resistance (Figure 5C; *r^2^*=0.68, *P*=0.003, Pearson correlation coefficient). No significant correlations were observed between cytotoxicity to macrophage and any of the other tested phenotypes (Figure 5 D-F).

**Figure 5.**
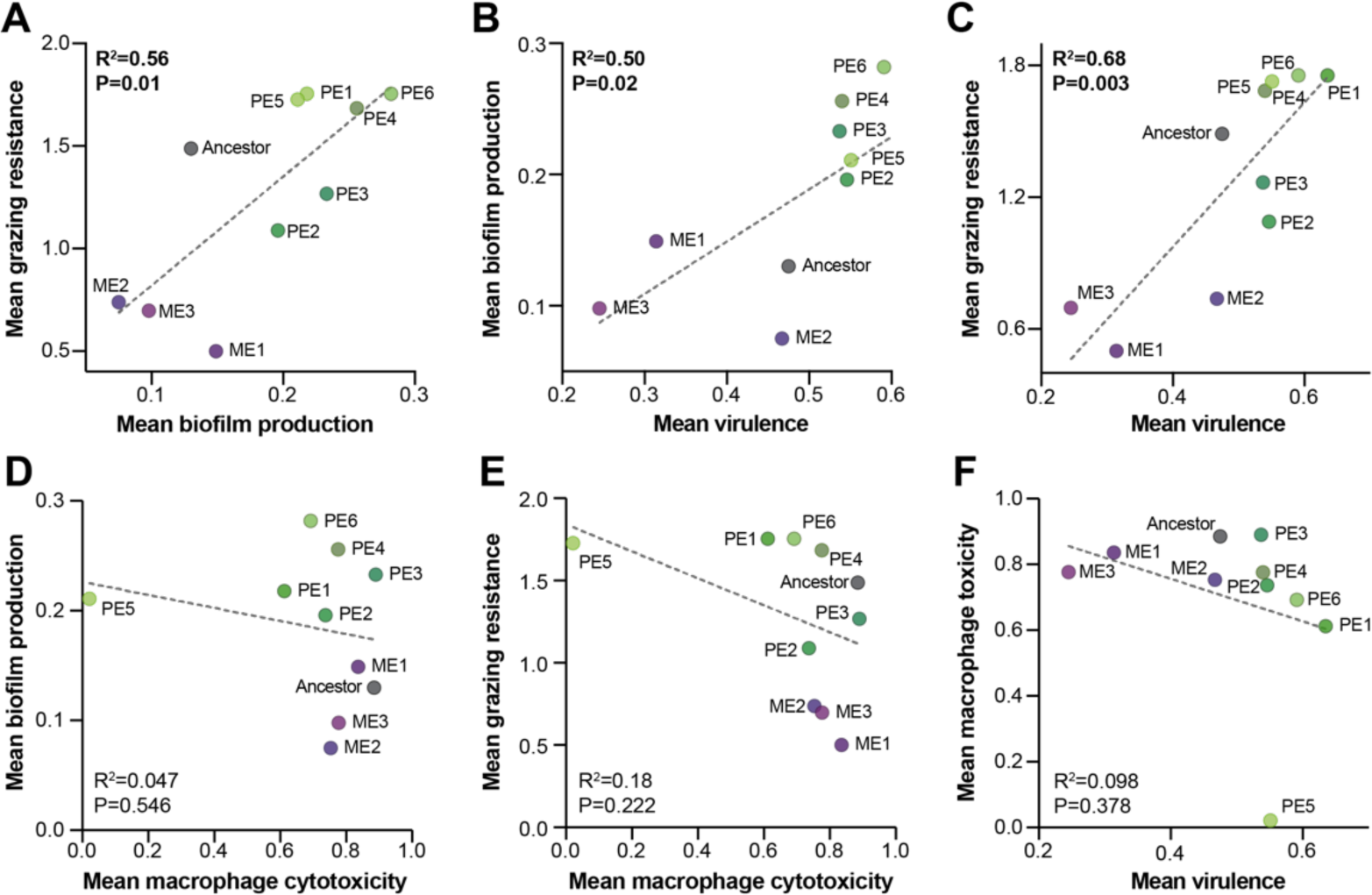
Correlations between mean. **A)** biofilm production and grazing resistance, **B)** biofilm production and virulence in bees, **C)** grazing resistance and virulence in bees, **D)** biofilm production and macrophage cytotoxicity, **E)** grazing resistance and macrophage cytotoxicity, and **F)** virulence in bees and macrophage cytotoxicity. Dashed lines represent a simple linear regression, and the correlation strength and significance were tested using the Pearson correlation coefficient.

## Discussion

Here we revealed that only 60 days of experimental evolution in the presence or absence of a predator resulted in overall consistent genotypic changes in *S. marcescens*, indicated by numerous parallel evolution events. The lower-than-expected number of nonsynonymous mutations coupled with the higher-than-expected number of parallel evolution events suggests that all evolved populations underwent both positive and purifying selection. We also found evidence of positive selection based on the presence of fixed mutations (sweeps) within the same genes, and sometimes in the same exact site, across populations. This is noteworthy as bottlenecks are inherent in experimental evolution and could result in the fixation of random mutations due to drift (66–68). The presence of fixed mutations in the same genes and positions strongly indicate that drift was not the main driver of the sweeps observed in our populations. Although parallel evolution events and sweeps suggest the genes impacted are adaptive, it is possible that some of the observed high frequency mutations are the result of hitchhiking.

Despite overall variation in the types of genes impacted, virtually all the fixed mutations within and across PEp and MEp populations occurred in genes that are part of the same predicted protein-protein interaction network. Within this protein-protein interaction network, several of the proteins are key components in gene regulatory systems. For example, BarA and UvrY make up a two-component regulatory system (49) and RcsB and RcsC are two of the three major components in the modified two component Rcs phosphorelay system (69), both of which have been shown to regulate the expression of numerous genes in *S. marcescens* (49, 69, 70) and have also been implicated in virulence mechanisms in multiple species of *Enterobacteriaceae* (52–56, 71–73). Thus, mutations in these genes likely have major impacts on multiple other genes by upregulating, downregulating, or stopping expression, resulting in strong selective pressure to maintain beneficial mutations and purge “deleterious” mutations. We hypothesize that mutations that were fixed within one or more genes in this pathway directed the evolutionary trajectories thereafter possibly due to epistatic interactions. However, further studies are needed to validate this hypothesis.

In order to link specific mutations with phenotypic characteristics, we randomly chose a single isolate from each population. It is important to note that for the MEp populations we only evaluated isolates from Lines 1-3 because when we originally started this experiment, we only evolved three ME populations. We later realized the importance of having more “control” populations and subsequently evolved three more MEp Lines, which were only analyzed at the population level in this study. However, we would like to point out that the three MEp Lines (MEp4-6) that were evolved over a year after MEp Lines 1-3 presented mutations in many of the same genes, including several of the fixed mutations that were observed in the first three MEp populations. This finding underlines the reproducibility of our results and eliminates the probability that contamination could explain the parallel evolution events observed across populations. The genotypic analysis of our three MEi isolates from Lines 1-3 and six PEi isolates (one from each evolved population) revealed that all isolates contained the fixed mutations present in their respective population. However, some isolates also contained mutations that were not fixed in the population. Moreover, only two isolates (PEi1 and PEi3) contained a single mutation, limiting our ability to directly correlate individual mutations with phenotypes. Based on our results, we predict that the sole mutations present in PEi1 and PEi3 in promoter regions upstream of the genes annotated as *pdeH* and a hypothetical protein, respectively, are responsible for the increased biofilm production observed in these two isolates. We also speculate that the large 66bp deletion in *rcsB* present in the MEi isolates contributed to the attenuated virulence in bees, as seen in MEi1 and MEi3 isolates. Furthermore, based on a previous study that demonstrated a role of the BarA-UvrY two component system in controlling the carbon storage regulatory system in *E. coli* (52), it is probable that the mutations we detected in *barA* and *uvrY* in the MEi and PEi isolates are the result of adaptation to the Neff media, which contains five percent glucose. We caution that these are only speculations that require further investigation to be validated.

The five phenotypes that we evaluated were growth in media, biofilm production, grazing resistance, virulence in honey bees, and cytotoxicity to murine macrophages. We found no change in population density (growth) of any of the isolates when compared to the ancestor in the three media types tested, but we did find changes in biofilm production. Specifically, all PEi isolates displayed increased biofilm production on average when compared to the ancestor, although PEi2 was not found to significantly produce more biofilm. The MEi isolates produced less biofilms than the PEi and ancestor on average, but individually none of them significantly produced less biofilm than the ancestor. As biofilm production is a natural defense mechanism against predators (74–76), increasing biofilm production could be beneficial in the constant presence of a predator, whereas it is likely not essential in a nutrient rich mono-culture. Although we did not see a significant increase in grazing resistance in any of the PEi isolates, we did observe a decrease in all three MEi isolates when compared to the ancestor. It is important to note that in order to maintain *T. thermophia* in co-culture with KZ19 for 24h at the start of the experiment we had to start with a ratio of only 10:1 (bacteria:protist) cell ratio, suggesting the ancestral strain already possessed mechanisms to resist grazing and/or survive phagocytosis. Thus, it is possible that in addition to the pressure to survive predation, competition for resources was a major force impacting the evolution of *S. marcescens* in the presence of *T. thermophila* in our experiments. In fact, although we found a strong positive correlation between biofilm production and grazing resistance, increased biofilm formation did appear to increase grazing resistance. Taken together, our results indicate that biofilm production promotes grazing resistance in *S. marcescens* KZ19 but also suggest that increasing biofilm production could be providing other functions in the presence of a predator aside from protection, such as the ability to better compete for resources (77, 78).

We found that both biofilm production and grazing resistance were positively correlated with virulence in a natural host (the honey bee). The correlation between grazing resistance and bee pathogenicity was particularly strong, especially given our sample size, providing support for the idea that predation can indirectly select for host virulence (5–8). Conversely, we found no correlation between cytotoxicity to murine macrophage cells (RAW 264.7) and biofilm production, grazing resistance, or virulence in bees. In fact, three of the PE isolates were significantly less toxic to macrophage after 2h of exposure, with PEi5 displaying virtually no cytotoxicity in this assay. Based on our time series experiment, we found that cytotoxicity to RAW 264.7 macrophages is highly time and density dependent. After 6h of co-culture, all isolates and the ancestral strain converged towards high cytotoxicity and had killed all macrophages in the culture. Other strains of *S. marcescens* have been shown to be highly toxic to macrophages (63–65), and our findings confirm this result for *S. marcescens* KZ19. However, the lack of correlation between macrophage toxicity and all the other phenotypes we tested suggests that the mutations present in our isolates have no impact on their ability to kill murine macrophage RAW 264.7 cells, and in some rare cases, might render them less toxic. These results indicate that, although evolution in the presence of a predator may increase bacterial virulence, the increase in virulence could be host specific, likely due to the diversity of mechanisms involved in virulence. This underlines the importance of testing the virulence of predator-evolved bacteria in multiple systems.

A few other studies have used experimental evolution to directly test how predation by *T. thermophila* impacts the virulence of *S. marcescens* (20, 21, 23). Contradictory to our findings, all these studies found that predation attenuated virulence. Some reasons for inconsistencies between our study and previous studies could be due to 1) the use of different *S. marcescens* strains with different life histories, 2) co-evolving *S. marcescens* and *T. thermophila* together (20, 21, 23) rather than preventing the evolution of the predator (this study), 3) using different hosts to test virulence, and 4) differences in the methods used to evaluate virulence. Virulence of the predator-evolved *S. marcescens* in previous studies was evaluated based on either testing entire populations (20) or individual isolates that were then pooled to evaluate virulence (21, 23). By testing a mixture of evolved isolates, it is likely that one will outcompete the others (and that the “winner” will be different depending on the environment).

Combining or comparing data from assays performed with different isolates could strongly impact the results since each isolate might genotypically and phenotypically differ (as we observed here) and be more or less fit depending on the context. Moreover, none of these studied evaluated the genotypes of the evolved populations, limiting their ability to fully evaluate how predation impacts the evolution of *S. marcescens*. Thus, we believe that the differences in experimental design between the previous predator *S. marcescens* evolution studies and ours likely explains the discrepancy in the results.

Fundamentally, our study demonstrates the efficacy of using experimental evolution to identify genes and pathways involved in virulence-associated phenotypes, even over short evolutionary timescales. Despite some variability in the evolutionary outcomes in our experiment, our results provide overall support for the hypotheses (i.e., CEH and the training grounds) that propose bacterial virulence is selected for or maintained by predation. Performing longer evolutionary experiments under different scenarios and with different opportunistic bacterial strains and species will provide a more accurate picture of the forces that drive the evolution of opportunistic pathogens and the mechanisms responsible for their virulence. Identifying virulence mechanisms and understanding how virulence evolves can help guide the development of more effective treatment strategies for combating bacterial infections.

## Materials and Methods

### Grazing confirmation

To confirm that *T. thermophila* grazes on *S. marcescens* KZ19, we utilized transconjugant KZ19 containing an E2 crimson fluorescent protein with spectinomycin resistance (38, 43). The transconjugant KZ19 strain was inoculated into 3mL of LB broth containing 180ug/mL spectinomycin and incubated at 30°C for 24h. After 24h, 1OD of the transconjugant KZ19 was resuspended in fresh Neff media and 1mL was pipetted into 3mL of Neff media containing *T. thermophila* and 180ug/mL spectinomycin. The culture was incubated for 24h at 30°C. One mL of the 24h co-culture was then centrifuged at 2000g for 10m to collect the *T. thermophila* cells, the supernatant containing bacteria was discarded, and the *T. thermophila* were resuspended in 1mL of axenic Neff media. The *T. thermophila* cells were then pipetted onto a microscope slide that was washed with 60% EtOH (which immobilizes the cells without immediately lysing them), covered with a glass cover slip, and imaged on a Keyence BZ-X700 Series All-In-One Fluorescence Microscope in brightfield and with a Cy5 filter.

### Experimental evolution

*Serratia marcescens* strain KZ19 was evolved either in media alone (media-evolved) or with the predator *T. thermophila* strain SB210 (predator-evolved). *T. thermophila* strain SB210 was purchased from the Tetrayhema Stock Center located at Cornell University. The media-evolved (ME) experiment included three replicate Lines and the predator-evolved (PE) experiment included six replicate Lines. First, *T. thermophila* and the ancestral *S. marcescens* KZ19 were cultured individually for 72h and 24h, respectively, at 30°C in Neff media. To start each evolved Line, 1OD of the overnight KZ19 culture was resuspended in fresh Neff media and 1mL was added to either 3mL of Neff media alone (MEp Lines) or 3mL Neff media containing *T. thermophila* culture (PEp Lines) at a ratio of 10:1 cells (bacteria:protist).The starting ratio of *S. marcescens*:*T. thermophila* was based on preliminary trials starting with various concentrations of *S. marcescens* and *T. thermophila* co-inoculated in Neff media to determine the ratio that would allow both organisms to remain in co-culture for 24h.

The 3mL experimental cultures were incubated at 30°C without shaking. Every 24h for 60 days the cultures were vortexed and 1% of each culture was sampled from the flasks and added to 3mL of either 1) fresh Neff media or 2) new axenic cultures of *T. thermophila* in Neff media. Every day, prior to passaging, the cultures were checked for *T. thermophila* viability and density via microscopy and the successful passaging and growth of *S. marcescens* presence was evaluated by plating on Luria-Bertani (LB) agar. Every day over the course of the 60-day evolution experiment, *T. thermophila* and *S. marcescens* were always detected. Although population size was not directly measured at every passage, based on daily plating (*S. marcescens*) and light microscopy (*T. thermophila*), we never observed a noticeable difference in population size. The number of generations reached for each *S. marcescens* evolved population over the course of 60 days is estimated to be approximately 396.

On day 60, six LB agar plates were streak inoculated from the culture flasks and incubated for 24h at 30°C. One bacterial colony was randomly selected from each of the six the PE populations (PE1-6) and three of the ME populations (ME1-3).Pure cultures of each isolate were created by inoculating in 3mL LB broth with 24h incubation at 30°C. Samples (200 μL from each pure culture) were then frozen at −80°C in 20% glycerol. These nine isolates were sequenced, analyzed, and used for phenotypic analysis (i.e., virulence in bees, growth in media, biofilm production, macrophage cytotoxicity, and grazing resistance).

### DNA Extraction and Sequencing

Cultures of the evolved populations (ME n=6, PE n=6), evolved isolates (ME n=3, PE n=6) and the ancestor were diluted to 1 OD. DNA extractions were performed using the Zymo Quick-DNA Fungal/Bacterial Miniprep Kit (D6005). DNA was prepped for sequencing using the Oxford Nanopore Rapid Barcoding Sequencing gDNA Kit (SQK-RBK004) for long read sequencing and the Illumina Nextera DNA Flex Library Prep Kit (20018704) for short read sequencing. The ancestral genome was sequenced via long (Oxford Nanopore Minion) and short (Illumina iSeq100) read technologies. The evolved populations and isolates were sequenced via short read technology on an Illumina iSeq100 with 2×150 paired end reads. See Table S1 for sequencing depth details.

### Genome Assembly and Analysis

The Illumina and Nanopore reads of the ancestral genome were then used to generate a hybrid assembly using hybridSPAdes v3.15.3 (79). Assembled scaffolds were annotated using Prokka v1.14.5 (80) on the Department of Energy Systems Biology Knowledgebase (KBase) platform (81). We also mapped all short reads of the ancestral strain back to the consensus ancestral genome using BreSeq v0.38.1 (45). Any mutations predicted by BreSeq when the ancestral reads were mapped to the ancestral consensus genome were considered sequencing or assembly issues in the ancestral genome and removed as candidate mutations in the evolved populations and isolates.

The sequencing reads of the evolved populations and isolates were trimmed using Trimmomatic v0.40 (82). The trimmed reads were then mapped to the newly assembled ancestral genome using BreSeq v0.37.0 (45) with a base-quality-cutoff PHRED score of 30 and minimum frequency cutoff of 0.05 to identify mutations. Promoters were predicted using iProEP (83). Functional characterization of genes (Figure 2-Table Supplement 2) was predicted using KOALA KEGG Orthology And Links Annotation (46, 84). Protein-protein interaction network prediction was determined using the STRING v12 database (48) using *S. marcescens*, *S. rubidaea* and *Y. enterocolitica* as reference organisms. Simulations to determine the percent mutations expected at random across intergenic, synonymous and non-synonymous positions and the probability of *gene-* and *site-specific* parallel mutations based on random expectation were conducted with a custom Python script: we conducted 10,000 independent simulations where one mutation was introduced at random under a Jukes & Cantor model in the reference genome of *S. marcescens* KZ19 (GCA_002915435.1). The annotations of the genome were used to infer whether the mutation was intergenic or not. Mutations affecting coding sequences were inferred as synonymous or non-synonymous by translating the gene *in silico* before and after introducing the mutation.

### Growth assay

Growth assays were performed in triplicate in both LB broth, fresh Neff media, and spent Neff media with *T. thermophila* filtered out (i.e., *T. thermophila* were grown in the media for 48h before filtering). Pure 1OD cultures of the evolved isolates and the ancestor were diluted to 10^-5^ and 5μL of the culture was pipetted into 200μL of media in a 96-well microplate. Absorbance detection was performed at an OD of 600nm at 30°C with shaking every hour for 24h on a BioTek Synergy™ 2 plate reader.

### Biofilm assay

Biofilm assays were performed in triplicate for each isolate. Pure 1OD cultures of the evolved isolates and the ancestor were diluted to 10^-5^ and 5μL of the culture was pipetted into 200μL of media in a 96-well microplate. The covered plate was incubated for 36h at 30°C without shaking. After 36h, the supernatant was poured off, the wells were washed twice with 200μL of sterile ddH2O, and the cells were stained with 150μL of 0.04% crystal violet for 10 minutes. After 10 minutes, the excess crystal violet was removed, the wells were washed again and stained once more for 10 minutes. The crystal violet was poured off and cells were solubilized with 150μL of 95% ethanol. OD readings were taken at absorbance reading 560nm on the BioTek Synergy™ 2 plate reader. A total of four assays were performed, each in triplicate.

### Grazing resistance assay

Grazing resistance assays were performed in triplicate for each isolate. Cultures of the evolved isolates and the ancestor were grown overnight, normalized the 1OD, and 1mL was added to a culture flask containing *T. thermophila* in 3mL Neff media (10:1 ration of bacteria to protist cells). *T. thermophila* alone in 3mL Neff media served as the negative control. Experimental culture flasks were incubated at 30°C for 48h. After 48h, the cultures were mixed thoroughly and 1mL was sampled and centrifuged at 5,000 rpm for 10 minutes. The supernatant was discarded, pellets were resuspended in 1mL 1X PBS, and the solution was filtered through 5.0 μm MilliporeSigma™ Millex™-SV Sterile PVDF syringe filters to remove the *T. thermophila* cells. OD readings were taken of filtrates at 600nm after blanking with the negative control filtrate. A total of two assays were performed, each in triplicate.

### Virulence assay

Pure 1OD cultures of the evolved isolates and the ancestor were pelleted via centrifugation and resuspended in a 1:1 sterile sugar syrup solution (SSS). Conventional adult honey bee workers (*Apis mellifera*) were sampled from a single hive located on the Gateway North Research Campus in Browns Summit, North Carolina. Bees were immobilized at 4°C, randomly distributed into groups, and exposed via the immersion method (38) to ∼10 μL of one of three treatments: 1) sterile SSS only, 2) the ancestor in sterile SSS, or 3) each evolved isolate in sterile SSS. Bees were kept in cup cages (20 bees/cup; 100 bees/treatment) under hive conditions: 35°C and 95% humidity. Mortality between the treatments was noted every day for 5 days. A total of three assays were performed with five replicates per assay equaling 300 bees tested in total per isolate. A survival curve (Kaplan-Meier) was created in GraphPad Prism v9.1.0.

### Macrophage cytotoxicity assay

Murine macrophage cells (Raw 264.7) were maintained in Dulbecco’s Modified Eagle’s Medium (DMEM, ATCC: 30-2002) with 10% Fetal Bovine Serum (FBS, ATCC: 30-2020) at 37°C with 5% CO2. After reaching 90% confluency, cells were harvested, resuspended in 500uL of DMEM media in a Corning™ Costar™ 24-well Clear TC-treated well plate (Fisher Scientific: 09-761-146), and incubated overnight at 37°C with 5% CO2. After overnight incubation, the old DMEM media was discarded, the macrophage cells were washed with 1X PBS, and 500μL of fresh DMEM media was added to each well. Overnight cultures of each bacterial isolate were normalized to 1 OD prior to the experiment and ten microliters of each 1 OD bacterial culture (MOI 10:1) was added into a separate well containing macrophage and incubated at 37°C with 5% CO2. For control wells (macrophage spontaneous release), 10μL of 1X PBS was added instead of bacteria. The cytotoxicity was measured after two hours using the CytoTox 96® Non-Radioactive Cytotoxicity Assay. In brief, two hours after co-culturing, the media from each well (including controls) was mixed two times and 60μL was pipetted into a 1.7mL tube and centrifuged at 10,000 rpm for two minutes to pellet bacterial cells. After centrifuging, 50μL of the supernatant was transferred into a new 96-well plate (Fisher Scientific: 07-200-90) to measure lactate dehydrogenase (LDH) release (OD490). For control wells, the macrophage lysis buffer release was measured by adding 50μL 10X lysis buffer to macrophage samples 45 minutes prior to LDH measurement, and the negative control was measured without adding lysis buffer. The percent cytotoxicity was calculated using the equation below. The ratio of 55/51 was used to correct the volume change caused by the lysis buffer. A single assay was performed with six replicates.

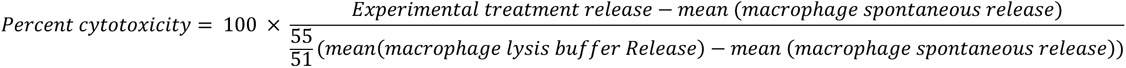

Since virtually all isolates were highly cytotoxic to macrophage after two hours at MOI of 10:1 cells (bacteria:macrophage), we performed a time series experiment to determine if macrophage cytotoxicity is time and density dependent. The Raw 264.7 macrophages and bacterial cultures were prepared as described above, except this time the assay started with a ratio of 1:10 (MOI) bacteria to macrophage cells and cytotoxicity via LDH release (CytoTox 96® Non-Radioactive Cytotoxicity Assay) was measured at five timepoints following co-culture (2h, 3h, 4h, 5h, and 6h). Cytotoxicity in this assay was based on optical density at OD490 rather than percent cytotoxicity. At 6h, all macrophages co-cultured with the ancestral and evolved isolates were dead as indicated by cell lysis observed under an inverted light microscope. One assay was performed in triplicate.

### Materials availability

All raw sequencing data has been deposited on NCBI under BioProject PRJNA838621.

## Acknowledgments

We would like to thank Dr. Carlos Goller for assistance with the Nanopore sequencing. We also want to thank Dr. Louis-Marie Bobay for helping with the simulations and providing constructive and helpful feedback on the manuscript. This work was supported by the National Science Foundation under grant DEB-2344788 (to K.R.), the National Institutes of Health under grant 7R01GM145747-02 (to K.R.), the University of North Carolina Greensboro (UNCG) College of Arts and Sciences Faculty First Award (to K.R.), and UNCG Department of Biology Graduate Student Support grants (to H.A.H and C.L.).

## Author Contributions

H.A.H., C.L. and M-J.L. performed the experiments and collected the data. K.R. designed and funded the research. All authors contributed data analysis and manuscript writing and editing.

## Supporting Material

**Figure S1.**
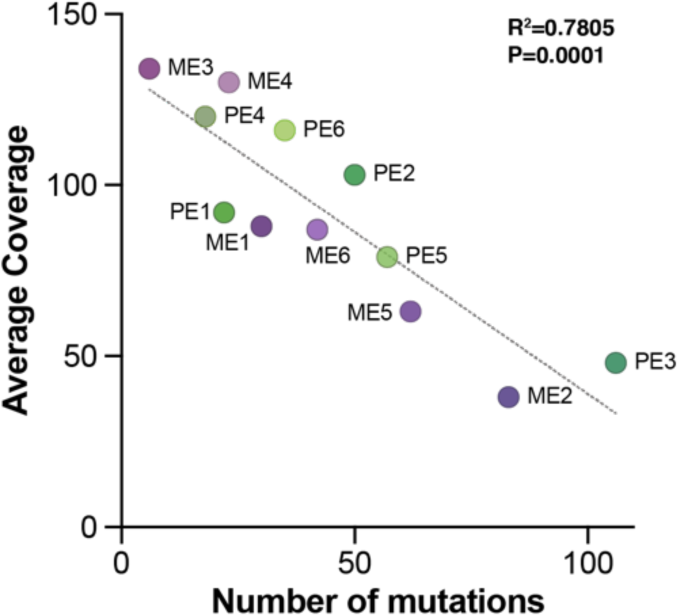
Comparison of the average genome coverage and the number of mutations detected. Dashed lines represent a simple linear regression, and the correlation strength and significance were tested using the Pearson correlation coefficient.

**Figure S2.**
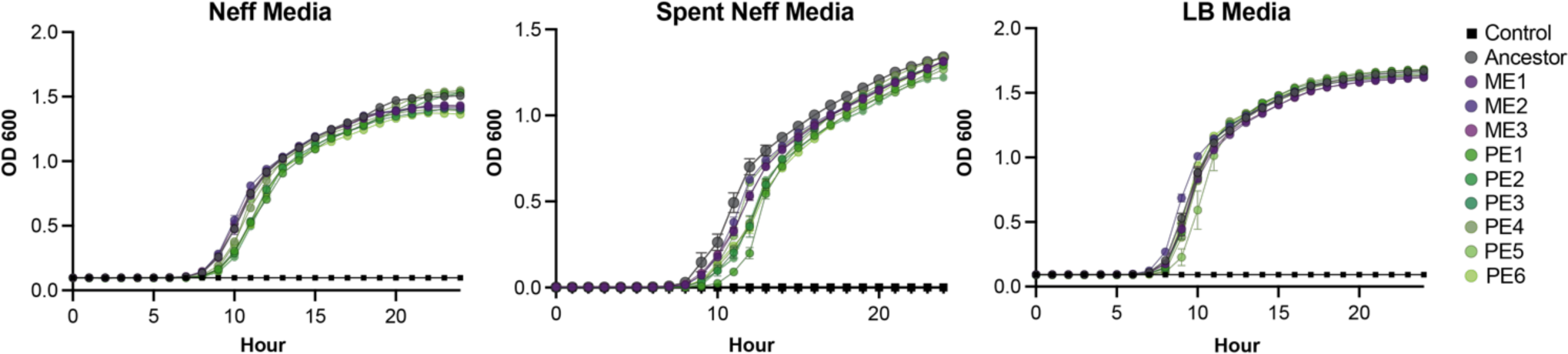
Mean growth rate of each variant based on 600nm OD readings every hour for 24h of ME (purple circles) and PE (green circles) isolates in LB broth (left) or fresh Neff media (right) compared to the ancestral strain (black circles). Significance was tested using Friedman One-Way ANOVA Repeated Measure Analysis with Dunnett’s multiple comparisons test.

**Figure S3.**
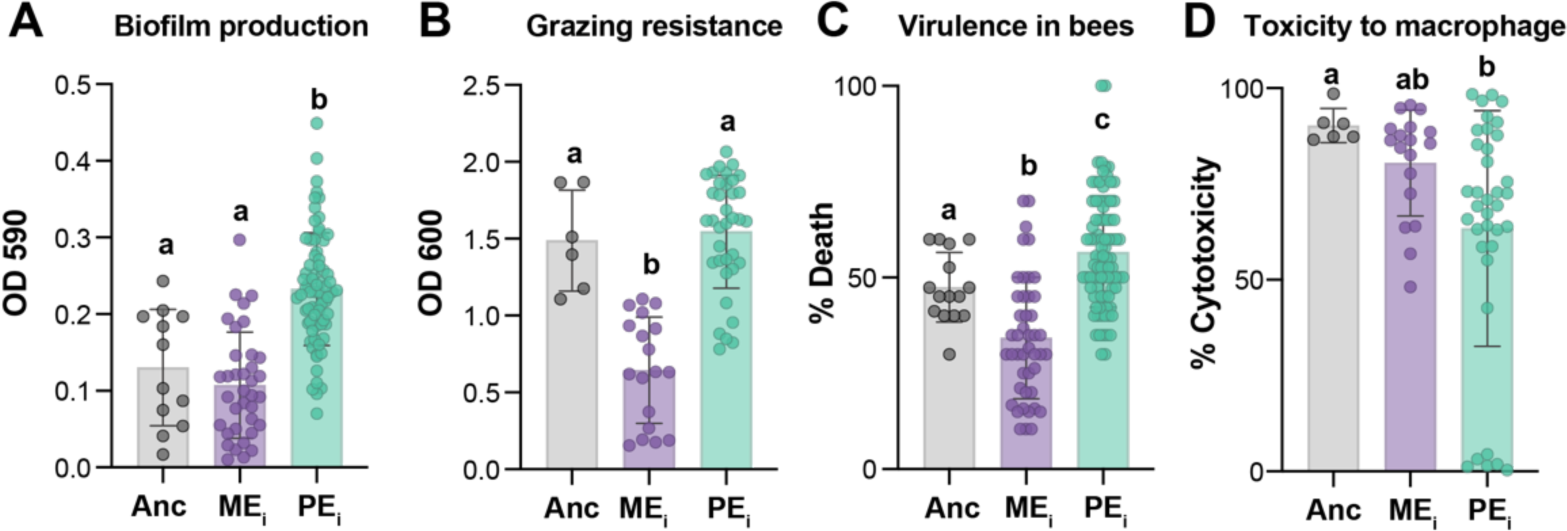
Mean A) biofilm production, B) grazing resistance, C) virulence in bees, and D) cytotoxicity to macrophage based on combining all ME and PE isolates. Significance was tested by comparing each evolved isolate to the others and the ancestor using one-way ANOVA with Dunnett’s multiple comparisons test.

**Figure S4.**
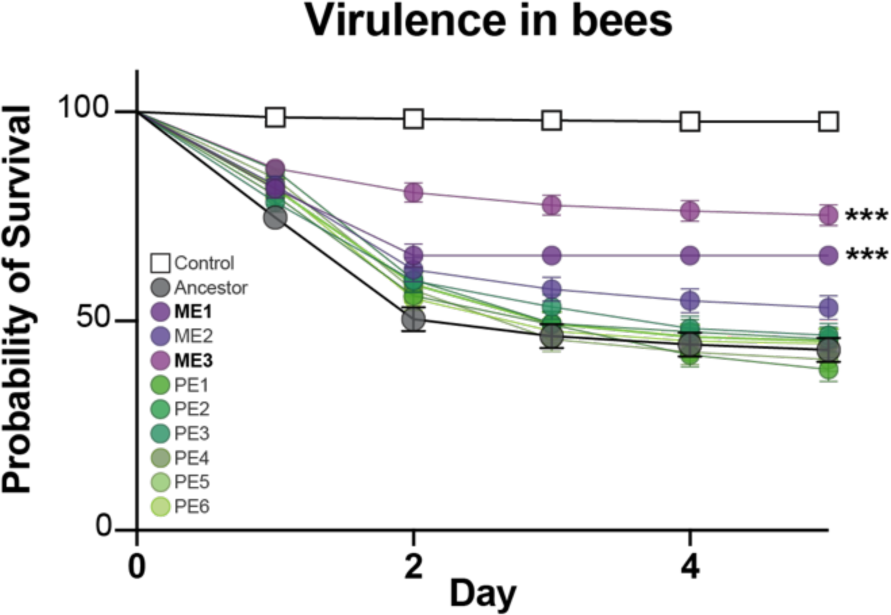
Kaplan-Meier curve showing the probability of survival of honey bees exposed to the ancestral strain (black and gray) and the **A)** ME isolates (purple) and **B)** PE isolates (green). Control bees were given sterile sugar water only (white squares). Mortality was recorded each day for 5 days. Graphs are based on three replicate assays per isolate (n=300 bees per isolate). * = P<0.05, *** = P<0.0005; Mantel-Cox Log-rank test with Bonferroni correction.

**Figure S5.**
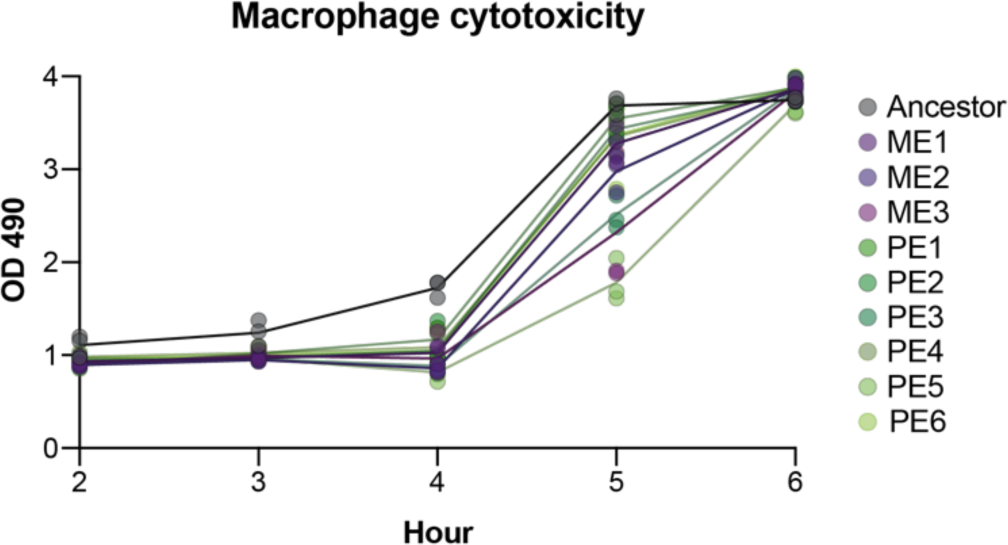
Cytotoxicity of MEi and PEi isolates and the ancestral strain to murine macrophage at an MOI of 1:10 (bacteria:macrophage) based on LDH release measured at OD490 after 2h, 3h, 4h, 5h, and 6h co-culture. The assay was done in triplicate. At 6h, death of all macrophage cells (lysis) was observed via microscopy by all evolved isolates as well as the ancestor.

**Table S1.** Total reads, number of mapped reads, and average genome coverage for each population and isolate.

**Table S2.** List of all the mutations identified in the evolved isolates and their frequency in the populations.

**Dataset S1.** List of all the mutations identified in the evolved populations and their frequencies.

