## Supplemental Figures for "Making a pathogen? Evaluating the impact of protist predation on the evolution of virulence in *Serratia marcescens*"

### Supporting Material

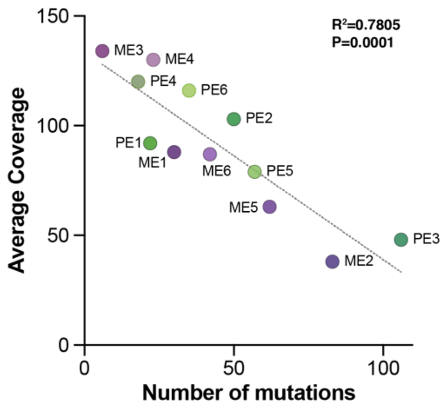


**Figure S1.** Comparison of the average genome coverage and the number of mutations detected. Dashed lines represent a simple linear regression, and the correlation strength and significance were tested using the Pearson correlation coefficient.

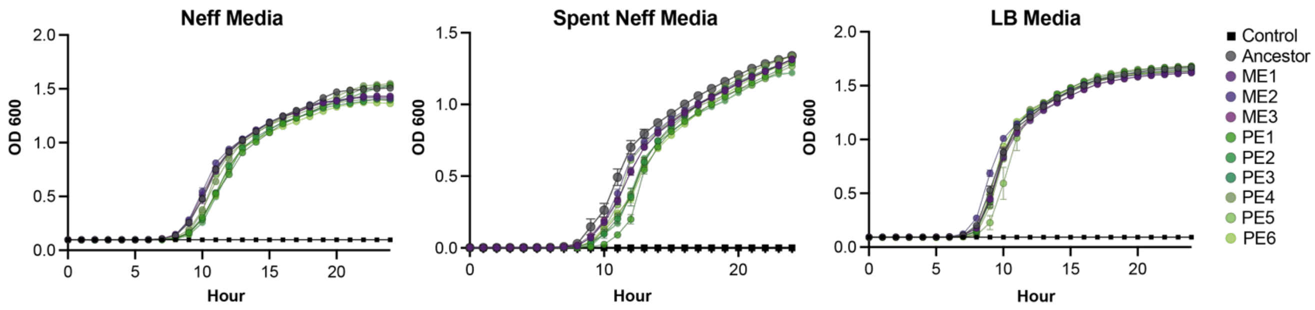


**Figure S2.** Mean growth rate of each variant based on 600nm OD readings every hour for 24h of ME (purple circles) and PE (green circles) isolates in LB broth (left) or fresh Neff media (right) compared to the ancestral strain (black circles). Significance was tested using Friedman One-Way ANOVA Repeated Measure Analysis with Dunnett's multiple comparisons test.

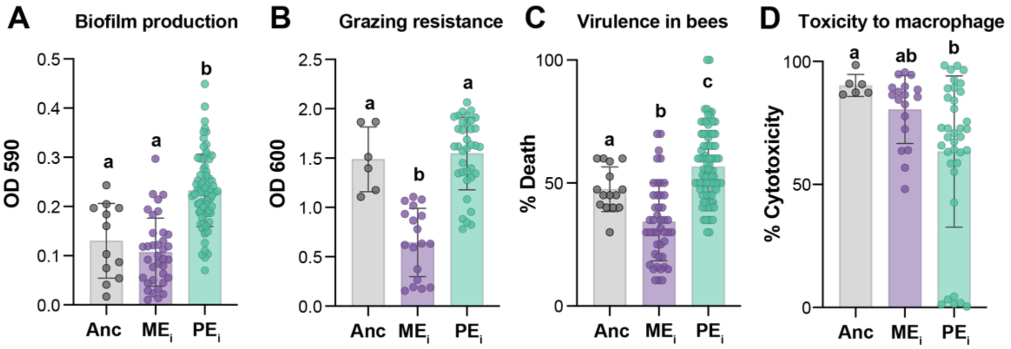

**Figure S3.** Mean A) biofilm production, B) grazing resistance, C) virulence in bees, and D) cytotoxicity to macrophage based on combining all ME and PE isolates. Significance was tested by comparing each evolved isolate to the others and the ancestor using one-way ANOVA with Dunnett's multiple comparisons test.

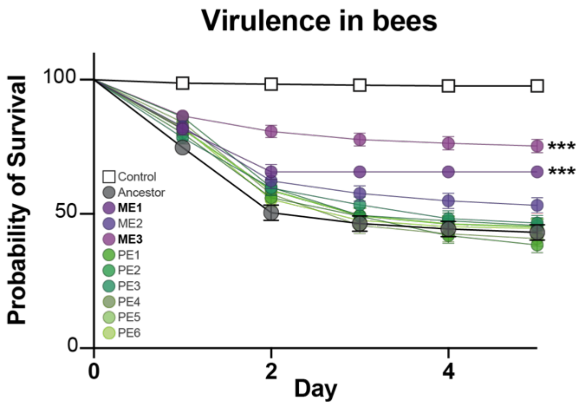


**Figure S4**. Kaplan-Meier curve showing the probability of survival of honey bees exposed to the ancestral strain (black and gray) and the **A)** ME isolates (purple) and **B)** PE isolates (green). Control bees were given sterile sugar water only (white squares). Mortality was recorded each day for 5 days. Graphs are based on three replicate assays per isolate (n=300 bees per isolate). * = P<0.05, *** = P<0.0005; Mantel-Cox Log-rank test with Bonferroni correction.

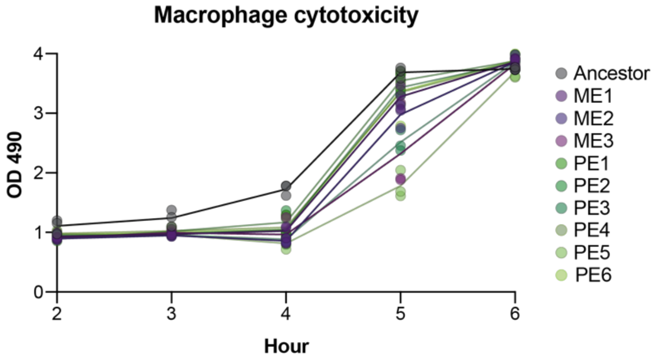


**Figure S5.** Cytotoxicity of MEi and PEi isolates and the ancestral strain to murine macrophage at an MOI of 1:10 (bacteria:macrophage) based on LDH release measured at OD490 after 2h, 3h, 4h, 5h, and 6h co-culture. The assay was done in triplicate. At 6h, death of all macrophage cells (lysis) was observed via microscopy by all evolved isolates as well as the ancestor.

**Table S1.** Total reads, number of mapped reads, and average genome coverage for each population and isolate.

**Table S2.** List of all the mutations identified in the evolved isolates and their frequency in the populations.

**Dataset S1.** List of all the mutations identified in the evolved populations and their frequencies.
